# Spontaneous alpha-band oscillations bias subjective contrast perception

**DOI:** 10.1101/2021.09.09.459569

**Authors:** Elio Balestrieri, Niko A. Busch

**Affiliations:** Institute of Psychology, University of Münster, Germany; Otto-Creutzfeldt-Center for Cognitive and Behavioral Neuroscience, University of Münster, Germany

**Author notes:** Corresponding author: Niko A. Busch, University of Münster, Institute of Psychology, Fliednerstrasse 21, 48149 Münster, Germany.

## Abstract

Perceptual decisions depend both on the features of the incoming stimulus and on the ongoing brain activity at the moment the stimulus is received. Specifically, trial-to-trial fluctuations in cortical excitability have been linked to fluctuations in the amplitude of pre-stimulus alpha oscillations (≈8-13 Hz), which are in turn are associated with fluctuations in subjects’ tendency to report the detection of a stimulus. It is currently unknown whether alpha oscillations bias post-perceptual decision making, or even bias subjective perception itself. To answer this question, we used a contrast discrimination task in which subjects reported which of two gratings – one in each hemifield – was perceived as having a stronger contrast. Our EEG analysis showed that subjective contrast was reduced for the stimulus in the hemifield represented in the hemisphere with relatively stronger pre-stimulus alpha amplitude, reflecting reduced cortical excitability. Furthermore, the strength of this spontaneous hemispheric lateralization was strongly correlated with the magnitude of individual subjects’ biases, suggesting that the spontaneous patterns of alpha lateralization play a role in explaining the intersubject variability in contrast perception. These results indicate that spontaneous fluctuations in cortical excitability, indicted by patterns of pre-stimulus alpha amplitude, affect perceptual decisions by altering the phenomenological perception of the visual world.

**Significance Statement:** Our moment to moment perception of the world is shaped by the features of the environment surrounding us, as much as by the constantly evolving states that characterize our brain activity. Previous research showed how the ongoing electrical activity of the brain can influence whether a stimulus has accessed conscious perception. However, evidence is currently missing on whether these electrical brain states can be associated to the subjective experience of a sensory input. Here we show that local changes in patterns of electrical brain activity preceding visual stimulation can bias our phenomenological perception. Importantly, we show that the strength of these variations can help explaining the great inter-individual variability in how we perceive the visual environment surrounding us.

## Introduction

Perceptual performance is not only determined by the external stimulus, but is also shaped by the ongoing internal brain state at the moment the stimulus is received. A specific type of ongoing brain signals are so-called alpha oscillations: rhythmic signals with a frequency of around 10 Hz, which are most dominant in visual cortical areas. Alpha oscillations reflect a state of reduced cortical excitability, indicated by an inverse relationship with single-unit firing rates (Chapeton et al., 2019; Haegens et al., 2011), multiunit activity (Bollimunta et al., 2008; Van Kerkoerle et al., 2014), or the fMRI BOLD signal (Goldman et al., 2002; Mayhew et al., 2013). In line with this inhibitory function, an increasing number of studies has demonstrated that the magnitude of alpha oscillations in the moment just before stimulus presentation affects the detection of near-threshold visual stimuli (Ergenoglu et al., 2004; Van Dijk et al., 2008; Lange et al., 2013). Specifically, strong alpha power induces a conservative detection bias, rendering observers less likely to report having seen a stimulus (Limbach and Corballis, 2016; Iemi et al., 2017; Iemi and Busch, 2018; Samaha et al., 2020a). While these studies have demonstrated the effect of pre-stimulus alpha oscillations on perceptual decision making, they raise a fundamental question about the relationship between spontaneous brain states and our subjective experience of the world: do fluctuations of cortical excitability reflect changes only in *decisions* about visual stimuli, or do they actually affect what the stimuli look like?

A similar question has been raised regarding spatial attention, which is known to improve accuracy in detection and discrimination tasks: is this improvement of objective performance associated with a change in the subjective appearance of the attended stimuli? To address this question, Carrasco et al. (2004) have developed a paradigm in which observers are presented with two Gabor patches, one on each side of fixation: a standard patch with a fixed contrast and a test patch whose contrast is randomly chosen from a range around the standard’s contrast. Observers report the patch with higher apparent contrast, resulting in a psychometric function indicating the probability of reporting the test patch as a function of the test’s relative contrast. Numerous studies have demonstrated that the point of subjective equality—the contrast at which the test stimulus appears subjectively similar to the standard—is shifted by both exogenous and voluntary attention, such that patches presented at attended locations appear subjectively more contrasted (see Carrasco and Barbot, 2019, for a review). Here, we adapted this paradigm to test if cortical excitability indicated by pre-stimulus alpha oscillations has a similar effect on contrast appearance.

To this end, we leveraged the fact that the amplitude of alpha oscillations is often lateralized across the left and right cortical hemispheres. Specifically, when spatial attention is cued to a lateral location, the distribution of alpha amplitude is shifted such that amplitude decreases in the contralateral relative to the ipsilateral hemisphere (Worden et al., 2000; Thut et al., 2006; Rihs et al., 2007), indicating greater excitability in the task-relevant hemisphere and greater inhibition in the task-irrelevant hemisphere (Jensen and Mazaheri, 2010). Accordingly, several studies have found that stronger cue-induced lateralization is beneficial for objective behavioral performance by speeding up responses for targets at attended locations (Gould et al., 2011) or by reducing the effect of distractors at unattended locations (Händel et al., 2011). Importantly, such lateralization can also occur spontaneously in the absence of external cues, driven only by internal fluctuations in the deployment of spatial attention, with similar effects on objective behavioral performance as cued shifts of attention (Bengson et al., 2014; Boncompte et al., 2016). Thus, our aim was to test whether subjective contrast appearance is amplified by such spontaneous pre-stimulus lateralization in the same way as by spatial cues. Furthermore, we sought to compare the topographical pattern of spontaneous lateralization to the well-known cue-induced lateralization of alpha oscillations.

To address this question, we used a paradigm similar to the one developed by Carrasco et al. (2004), but without attentional cues. Thus, instead of comparing the point of subjective equality between cued vs. non-cued stimuli, we compared the spontaneous pre-stimulus alpha lateralization between trials when the left vs. right stimulus was perceived as most contrasted.

In line with our expectations, based on the well-known inhibitory effect of alpha oscillations, we found that stimuli preceded by reduced contralateral compared to ipsilateral amplitude appeared as stronger contrasted. For comparison, we included an additional task with symbolic attentional cues, which induced typical lateralization with reduced alpha amplitude contralateral to the cued hemifield. The topographical pattern of cue-induced lateralization was highly similar to that of the pre-stimulus lateralization effect on subjective contrast appearance.

Using a logistic regression approach, we were able to determine the spatiotemporal evolution of the spontaneous pre-stimulus lateralized patterns that biased subjective appearance on a trial by trial basis, alongside with a non lateralized effect of spontaneous pre-stimulus alpha that impacted on the quality of perceptual representation. Finally, we found that the strength of the spontaneous lateralization patterns significantly correlated with the PSEs across participants, strengthening the claim that local shifts in pre-stimulus alpha amplitude bias appearance.

## Results

Human observers (N=40) performed a contrast discrimination task (Figure 1). Two Gabor patches were presented on each trial, one in the lower left and another in the lower right visual field (VF). One of the patches had a fixed contrast (standard), while the contrast of the other patch (test) was selected from a range around the standard’s contrast. Observers reported which of the patches appeared to have the highest contrast, resulting in a psychometric function (PMF) describing the probability of reporting the test patch as a function of the test’s relative contrast. Test contrasts were determined trial-by-trial by an adaptive Bayesian algorithm, which selected the test contrast that would be most efficient for estimating the PMF based on the data collected so far (see Methods). In a separate cued orientation discrimination, from hereafter named “attention localizer”, the Gabors were preceded by a symbolic cue indicating the location of the to-be-attended and to-be-reported target. Observers reported the orientation of the target patch (+45° or −45°).

**Figure 1:**
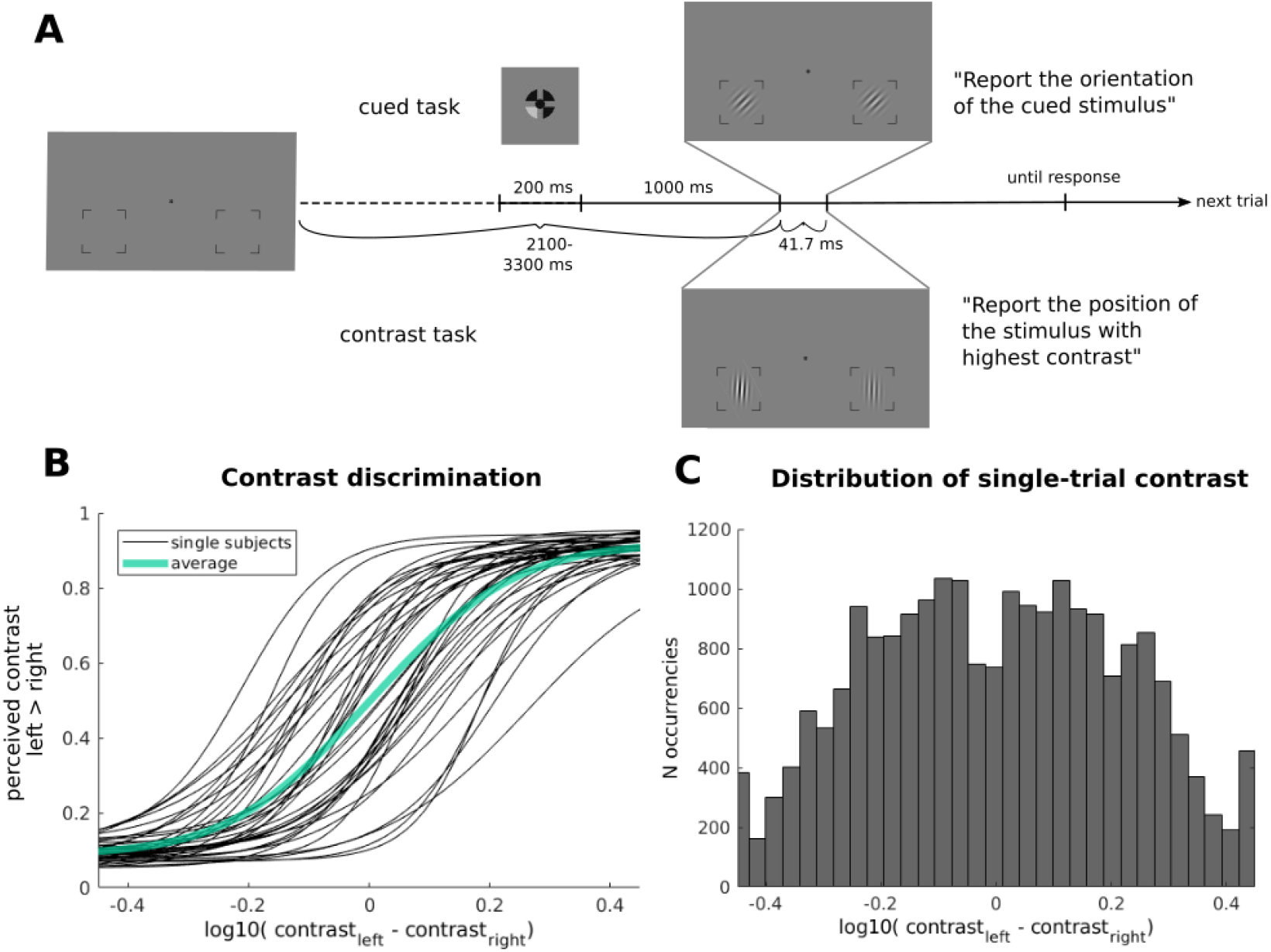
**A)** Illustration of the experimental paradigm. Each attention localizer trial started with a symbolic cue indicating the position to attend, followed by the simultaneous presentation in the left and in the right low visual field of two Gabor patches, both at the same contrast level, each of which could have an orientation of ±45°. In the contrast discrimination task no cue was provided, but only the two Gabor patches, this time both with vertical orientation. One of them always had a stable contrast throughout the experiment, whereas the contrast of the other was allowed to change. In the attention localizer task the participants had to report the orientation of the cued stimulus, whereas in the contrast discrimination task they had to report which stimulus they perceived as “more contrasted”. **B)** PMFs estimated for the single subjects and the average PMF as *P_resp_* toward the left stimulus as a function of the contrast difference between *left* and *right*. **C)** Distribution of contrast values probed across the whole set of participants.

### Behavioral Results

Participants performed well in both tasks, achieving a performance of .748 ± .091 (mean ± SD) in the contrast discrimination task (averaged across all test contrasts) and .978 ± .03 in the attention localizer task.

For the contrast discrimination task, we used data from all trials to re-estimate parameters of a PMF, namely the PSE (Point of Subjective Equality) and slope, describing for each observer the probability of choosing the left patch as the most contrasted as a function of the contrast difference between the left and right patch (Fig. 1B). While individual observers showed a general preference for choosing patches either in the left or right hemifield, these preferences were balanced across observers. Thus, the resulting average PMF was almost perfectly centered on a contrast difference of zero, indicating no systematic bias towards either hemifield.

The distribution of absolute contrast differences was balanced across hemifields, such that trials with stronger contrast in the left hemifield were stronger by the same magnitude as trials with stronger contrast in the right hemifield (Fig. 1C). This symmetry was confirmed by a two-tailed t-test comparing absolute contrast differences between trials with stronger contrast in the left vs. right hemifield (*t*_(20903)_ = −.114, *p* = .909). As a result of the adaptive Bayesian procedure, the majority of trials featured small, but non-zero contrast differences (Fig. 1C) around the PMF’s midpoint. For these trials, subjective contrast appearance was highly variable and not fully determined by the objective contrast difference, indicating high perceptual uncertainty. This allowed us to test whether contrast appearance was influenced by lateralized pre-stimulus oscillatory amplitude.

### Effect of spontaneous pre-stimulus lateralization on contrast appearance, and its relationship with cue-induced alpha lateralization

We analyzed pre-stimulus lateralization using a time-frequency analysis across the 1 second pre-stimulus window and a broad range of frequencies (3–40 Hz), and compared lateralization between trials in which the left vs. right patch was reported as more contrasted. This comparison is a valid test of subjective contrast appearance even though it disregards the objective contrast difference between left and right patch, and thus is likely to include trials on which the report was determined by substantial objective contrast differences. First, as shown above, the majority of trials featured only subtle contrast differences, and these differences were of the same magnitude in the left and right hemifield (Fig. 1C). Second, we only analyzed the pre-stimulus time range, uncontaminated by the neural post-stimulus response (see Methods). Thus, any pre-stimulus differences between post-stimulus left-reports and right-reports are necessarily independent of objective stimulus contrast.

At each frequency, we computed lateralization as the amplitude difference between a group of left minus right occipital channels, and compared this lateralization between trials in which the left vs. right patch was reported as more contrasted. This comparison showed a positive difference in lateralization between left and right reports, starting approximately 200 ms before stimulus onset (Fig 2 A), indicating that in both conditions, alpha amplitude was reduced at channels contralateral compared to ipsilateral to the patch reported as more contrasted.

**Figure 2:**
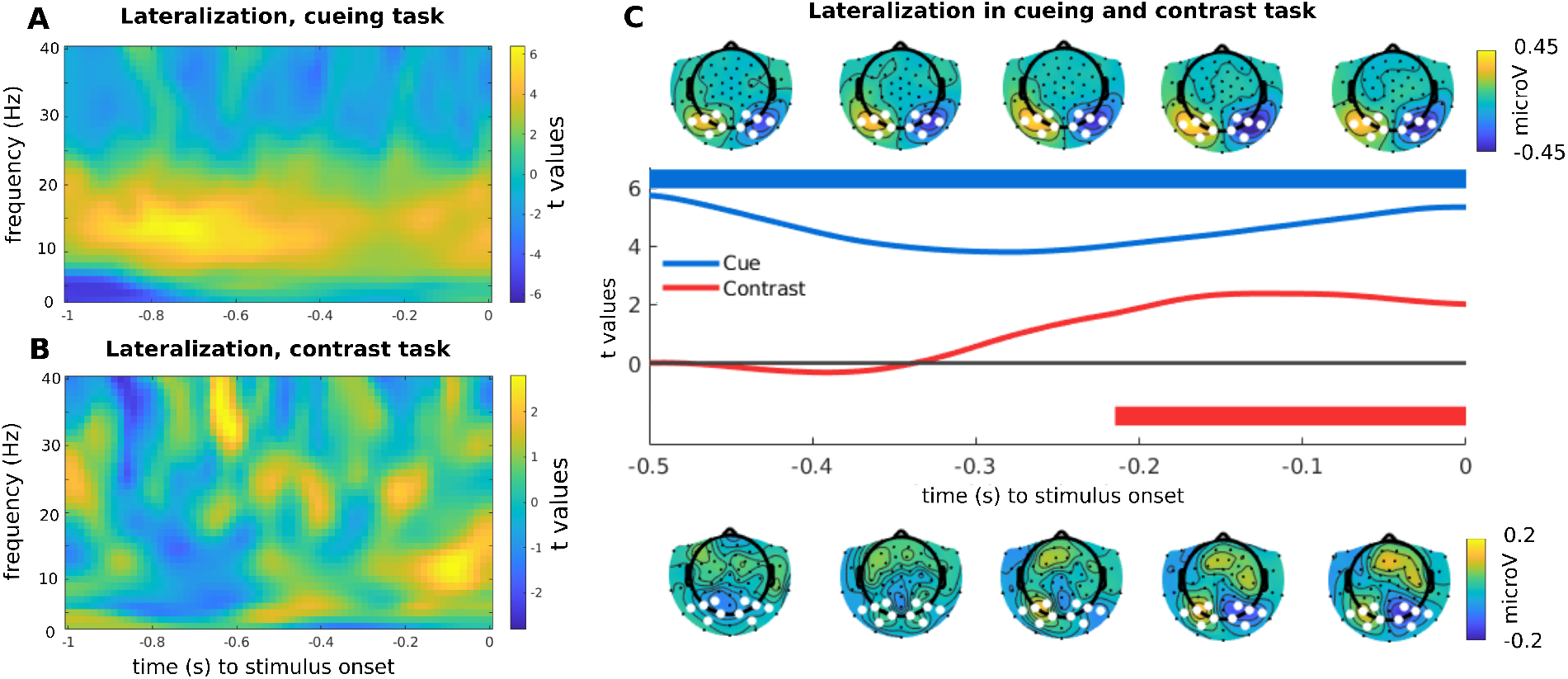
**A)**. Time frequency difference over a broad window of 1000 ms pre-stimulus between left vs right occipital channels, in the comparison between left cue vs right cue trials. It is possible to notice a great increase in amplitude spanning the whole pre-stimulus window in a wide range of frequencies in the alpha-beta band. **B)**. Same analysis, but now on the difference between high contrast perceived to the left vs right. The effect of interest here spans a much reduced time window, and frequencies. **C)** Inter-channel difference in alpha amplitude, cueing vs contrast tasks. The upper topographies refer to the cue, the lower to the contrast response direction. Each topography averages the amplitude scalp distribution in the alpha range (8-13 Hz) over a time window of 100 ms, for the 500 ms preceding stimulus onset. Highlighted channels are the ones used for the cluster permutation test. Blue refers to cue condition (left - right), Red refers to response direction in contrast condition (left - right). The upper and lower bars show the portion of the significant cluster over time (difference from 0), color coded as before.

To confirm this result statistically, we band-pass filtered the data in the alpha-band, obtained instantaneous amplitude with a Hilbert transform, and computed again the difference between left and right occipital channels in the contrast between left and right responses, as described above. Similar to the time-frequency analysis, this contrast, whose sign indicates the direction of lateralization, was positive in the interval just before stimulus onset. A non-parametric cluster permutation test yielded a cluster of significant non-zero lateralization (*P* = .025) starting at −214 ms before stimulus onset (red horizontal bar in Fig. 2C). This result confirms that alpha amplitude was significantly reduced in the cortical hemisphere contralateral to the patch reported as more contrasted. Note that this analysis was conducted on data from which any post-stimulus data points were removed (see Methods). Thus, while this pre-stimulus effect occurred just before stimulus onset, it cannot be explained by temporal smearing of post-stimulus signals into the pre-stimulus time range. Cue-induced lateralization in the attention localizer task showed a similar, but stronger and more sustained effect (cluster *P* < .001).

If, as hypothesized, spontaneous patterns of alpha lateralization are related to a change in subjective perception analogous to changes produced by spatial attention shifts, it is reasonable to expect higher similarity between spontaneous and cue-induced alpha lateralization under maximal perceptual uncertainty, i.e. for contrast values around the PMF’s midpoint. To test this prediction, for each participant, we split the trials in high and low uncertainty in the contrast discrimination task based on the individual PMF (see Fig. 3B and Methods) and computed for each set of trials lateralization as the difference between left and right choices, as described before. We then computed the similarity matrix between the pattern elicited by the attention localizer task and the pre-stimulus pattern associated to the response both in high and low uncertainty.

**Figure 3:**
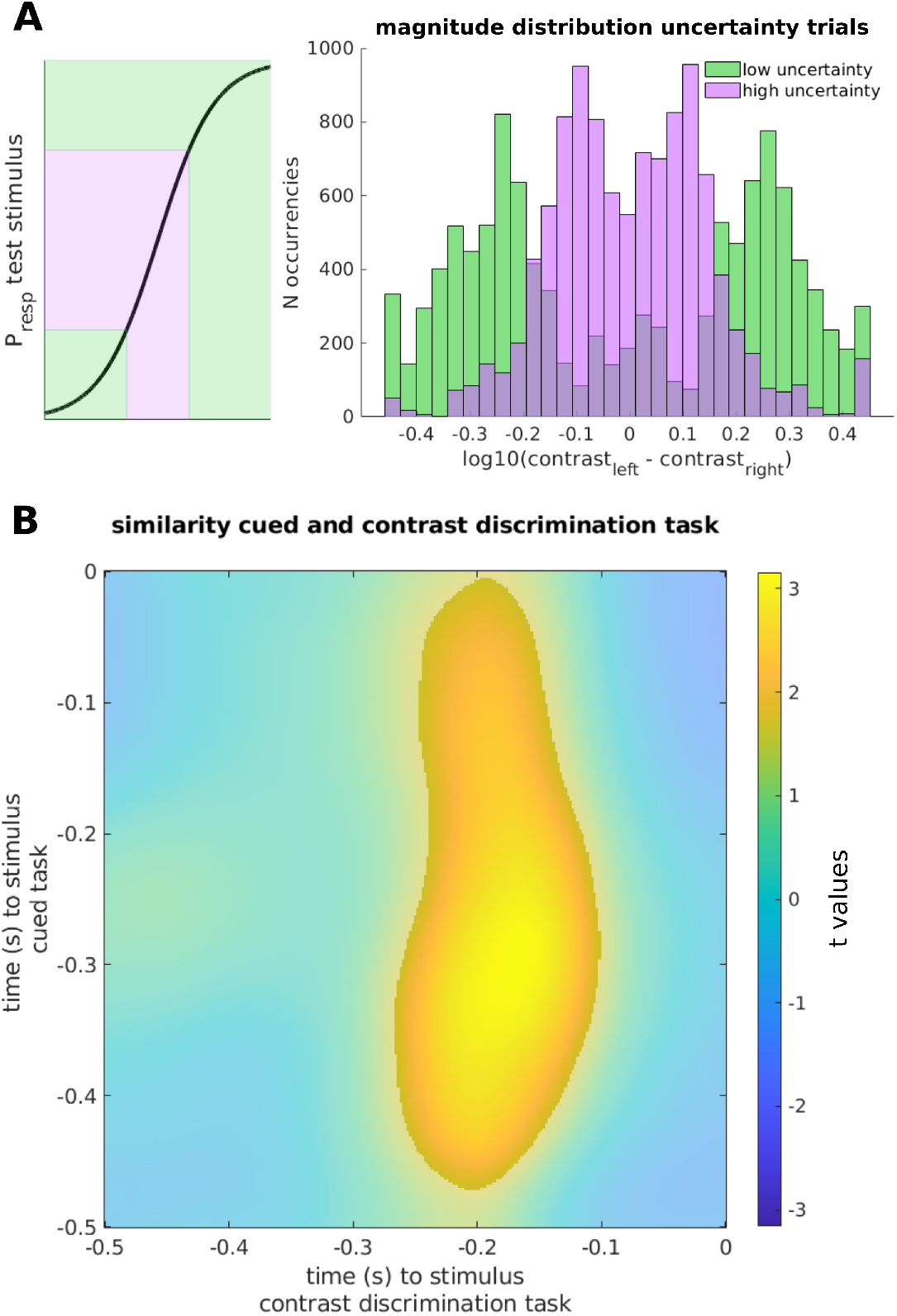
Similarity between cue induced and spontaneous lateralization, in the comparison between high and low perceptual uncertainty. **A)** The definition of trials with high uncertainty was performed by selecting a range of ± .25 around the point of subjective equality, defined as the point where the probability of response for either of the hemifields is .5. This resulted, across all participants, in two distinct sets of trials (color coded distributions in right inset). **B)** Similarity matrix between the cue-induced and the spontaneous lateralization, in the comparison between high and perceptual uncertainty as defined in **A**. The comparison yields a significant cluster of similarity around 200 ms pre-stimulus, indicating that the pattern of spontaneous lateralization is more congruent to the one induced by the cue when there is higher perceptual uncertainty.

This comparison of the similarity matrices in these two different perceptual uncertainty conditions yielded a significant cluster (*P* = .037), centered around 200 ms pre-stimulus for the contrast discrimination task but spanning almost the whole time window for the attention localizer task (Fig.3C). This result implies that under high perceptual uncertainty the appearance of one stimulus as more contrasted is preceded by a short-lived lateralization of alpha amplitude, whose topographical pattern is similar to the conventional cued-induced lateralization pattern.

### The influence of pre-stimulus alpha amplitude on trial by trial perceptual decision making

While the previous analyses compared pre-stimulus lateralization between conditions defined by the subject’s post-stimulus choices in the contrast discrimination task, the hypothesized chain of causality is actually the opposite: pre-stimulus brain states may bias post-stimulus choices. However, the analysis of effects of pre-stimulus lateralization on decisions must account for the additional effect of objective contrast differences. Therefore, to analyze the effect of trial-by-trial variations in pre-stimulus alpha-band amplitude on perceptual decisions while controlling for objective contrast differences, we used a logistic regression analysis, which modeled the probability of reporting the left stimulus as more contrasted on a given trial based on four regressors: a constant intercept reflecting the subject’s individual preference for (or against) reporting the left side, single-trial pre-stimulus alpha amplitudes, single-trial contrast differences between left and right stimulus, and the interaction between amplitude and contrast differences. This model was applied independently for each participant, channel, and time point.

Fig. 4A shows topographies indicating for each channel the effect of single-trial pre-stimulus alpha-band amplitude on the probability of reporting the left stimulus as more contrasted. Thus, positive values indicate an alpha-induced bias for left reports. These effects are shown across several time windows leading up to stimulus onset, illustrating a significant cluster (*P* = .03) that evolves from a more central topography to a left posterior topography just before stimulus onset. This effect indicates that strong alpha amplitude in the left posterior hemisphere biased subsequent contrast perception against stimuli presented in the contralateral hemifield, or towards stimuli in the ipsilateral hemifield independently of objective contrast differences, thereby confirming the results of our previous analyses. To further illustrate how this biasing effect plays out for different objective contrast differences, we grouped trials into five bins according to their contrast difference. Then for each subject, we selected the channel with the highest beta score, indicating that strong alpha amplitude at this channel was most predictive for a leftward response, and the channel with the lowest beta score. We then inverted the model to compute the probability of a leftward response for each of the five contrast bins, separately for trials with strong or weak amplitude at each of the two channels of interest. We then computed for each contrast bin and each channel the difference in probability for a leftward response on strong minus weak amplitude trials, and in turn subtracted this difference at the channel with highest beta score from the channel with lowest beta score. Thus, the result of this double-subtraction indicates which objective contrast differences were most affected by the biasing effect of pre-stimulus alpha amplitude. As shown in Figure 4C, this effect was strongest for the central contrast bin comprising contrast differences around the point of subjective equality, i.e. trials with highest perceptual uncertainty.

Fig. 4B shows topographies indicating for each channel the interaction between pre-stimulus amplitude and objective contrast difference on the probability of a leftward response, illustrating a significant, non-lateralized negative cluster (*P* = .024) throughout the 500 ms pre-stimulus interval concentrated at parieto-occipital channels, extending also to frontal channels. To further illustrate the interaction of alpha amplitude and objective contrast differences, we again grouped trials into five bins according to their contrast difference, separately for trials with strong and weak amplitude at each subject’s channel and time point featuring the highest beta score. As shown in Figure 4C, the probability of a leftward response generally increased with increasing objective contrast of the left stimulus, thus reflecting the psychometric function (see 1D). However, the slope of this relationship was steeper for trials with weak alpha amplitude, indicating that strong, non-lateralized alpha amplitude generally impaired the accuracy of perceptual decisions. In sum, our single trial regression approach allowed us to uncover two distinct effects of prestimulus alpha amplitude on perceptual decision making: a bias effect whereby strong lateralized alpha amplitude just before stimulus onset reduced the perceived contrast in the contralateral hemifield, and an inhibitory effect whereby strong non-lateralized alpha amplitude impaired perceptual accuracy.

**Figure 4:**
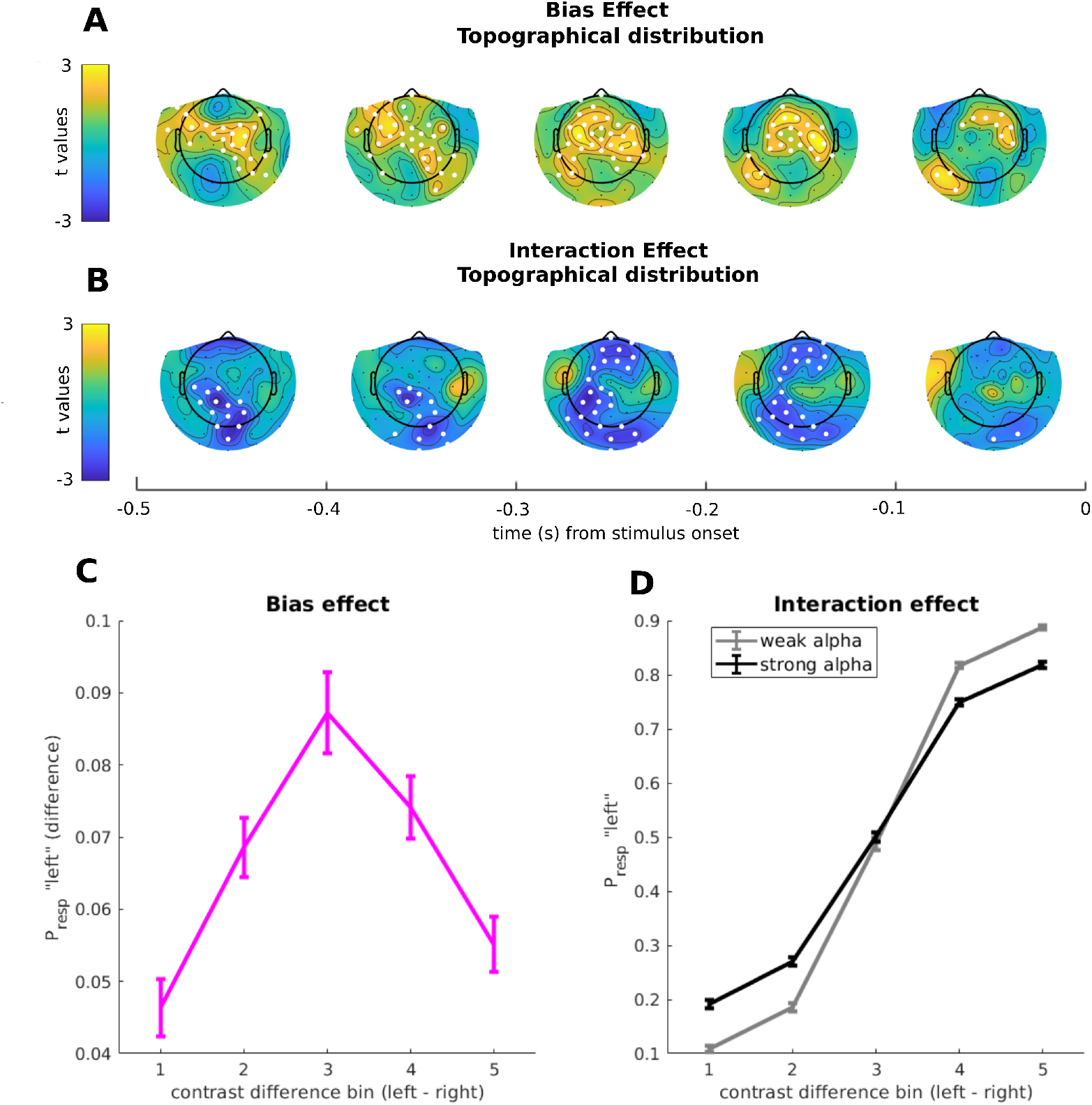
Single trials logistic regression. **A)** Spatiotemporal distribution of beta scores representing the effect of single-trial pre-stimulus alpha-band amplitude on the probability of reporting the left stimulus as more contrasted: from a widespread topography in the early time windows, it is possible to observe the build up of a lateralized pattern in the 200 ms preceding stimulus onset. Highlighted in white are the channels participating to the cluster formation. **B)** Spatiotemporal distribution of beta scores representing the interaction between alpha amplitude and physical contrast of the stimuli. The increase of alpha amplitude mostly located in occipito-parietal, and, for a short transient, in frontal areas, negatively interacts with the physical contrast of the stimuli. Highlighted in white are the channels participating to the cluster formation. **C)** Representation of the bias effect. We first binned the trials in 5 different contrast levels, and we inverted logistic regression model for each bin, for the couple of channels with the highest (positive) and lowest (negative) beta scores. The subtraction of the probabilities of response toward the left hemifield shows a more pronounced biasing effect around the central contrast bin, which comprises the point of subjective equality, indicating maximal biasing effect under perceptual uncertainty. **D)** Representation of the interaction effect. With the same 5 bins used before, we selected the channel with the highest beta score for the interaction between physical contrast difference and alpha amplitude, and further binned single trials in strong and weak alpha amplitude. Then we inverted the logistic regression for each contrast and alpha amplitude bin: this representation shows that strong, non-lateralized alpha amplitude generally impaired the accuracy of perceptual decisions.

### The strength of pre-stimulus alpha lateralization correlates with individual PSE

The previous analyses demonstrated that within-subject, trial-by-trial fluctuations of pre-stimulus alpha-band lateralization have a biasing effect on contrast perception in the left and right hemifield. At the same time, we also observed strong *across-subjects* variability in the general preference to report either hemifield as more contrasted (see Figure 1B), possibly indicating inter-individual variability in subjective contrast perception in the two hemifields. Thus, we tested whether subjects’ individual general preferences for either hemifield is related to the strength of their individual lateralization-induced biases.

To this end, we quantified the strength of each subjects’ alpha-band lateralization as the difference in amplitude between left and right occipital channels, and their lateralization-induced biases as the difference in pre-stimulus lateralization between left and right responses in the contrast discrimination task. Thus, positive values of this metric indicate that reporting a particular hemifield as more contrasted is preceded by stronger amplitude at ipsilateral compared to contralateral electrodes (i.e. the pattern shown in Figure 2A and Figure 4A), while negative values indicate a reversed pattern. We expected a positive correlation between subjects’ prestimulus lateralization-induced biases and their individual PSEs, meaning that subjects with a strong association between left lateralization and left responses in the contrast discrimination task would also show a general preference for the left hemifield, and vice versa. Indeed, we found a significant cluster (Fig. 5A, *P* = .003) spanning a great portion of the pre-stimulus window. The highest peak of the correlation, at ≈ 379 ms before stimulus (*r* = .391, *p* = .006) is shown in Fig. 5B.

This result demonstrates that participants with an overall preference to report the left stimulus as more contrasted also showed a congruent lateralization pattern in the difference between left vs right responses, and vice versa, suggesting that the large inter-individual variability in subjective contrast perception is partially rooted in the same patterns of cortical pre-stimulus excitation and inhibition that also bias trial-by-trial decision making.

**Figure 5:**
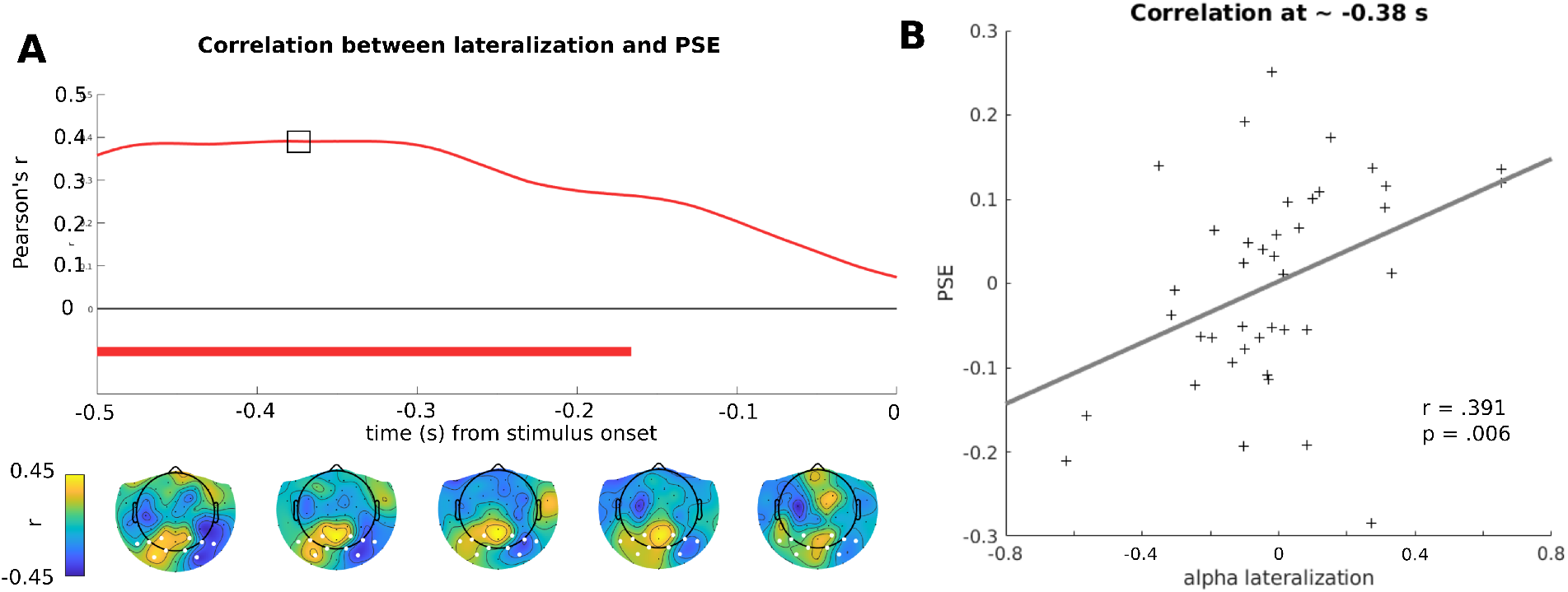
**A** (***Upper inset***) Timecourse of the correlation between spontaneous alpha lateralization and individual PSEs. This indicates that the lateralization pattern associated to enhanced contrast on the left hemifield against the right is correlated with the strength of the individual biases (i.e. the shift from 0 of the PSE). The small square indicates the position over the timecourse of the peak correlation, shown in **B**. (***Lower inset***) topographical distribution over time of *r* values obtained by correlating the pattern of spontaneous lateralization and with individual PSEs. The channels used for computing alpha lateralization, the same in 2C, are highlighted in white. **B)** Peak correlation between alpha lateralization and individual PSEs, at 380 ms before stimulus onset.

## Discussion

Do spontaneous fluctuations in alpha amplitude alter perceived contrast? To answer this question, we used a contrast discrimination task in which two Gabor patches were presented in the left and right hemifield, and participants reported the stimulus with the highest contrast (Carrasco, 2011). Based on the finding that strong alpha amplitude is associated with reduced cortical excitability (Romei et al., 2008; Bollimunta et al., 2008; Haegens et al., 2011), we expected that perceived contrast of the stimulus represented in the more excitable cortical hemisphere would be amplified. Across several analyses, we confirmed this relationship between contrast appearance and pre-stimulus alpha amplitude lateralization just before stimulus onset. Specifically, stimuli contralateral to the hemisphere with weaker pre-stimulus alpha amplitude were more likely to be reported as more contrasted (Figure 2B, 2C, 4).

Numerous studies have demonstrated that the amplitude of alpha-band oscillations is not only related to physiological excitability, but also to behavioral performance (Ergenoglu et al., 2004; Van Dijk et al., 2008): in detection tasks, weak alpha amplitude, indicating increased cortical excitability, is associated with a more liberal detection criterion (Iemi et al., 2017; Iemi and Busch, 2018), higher confidence (Samaha et al., 2017), and higher subjective stimulus visibility (Benwell et al., 2017), but without a change in accuracy. At present, it is still debated whether these effects indicate a change at the level of subjective perception of the visual world, or a change at the subsequent decision level (Kloosterman et al., 2019; Samaha et al., 2020a). In a simple detection task requiring a target presence/absence judgment, a bias at the perceptual level would imply that any stimulus (be it target or noise) looks more target-like, while a bias at the decision level would imply that participants feel more inclined to report target presence. In the comparative contrast discrimination task used in the present study, however, a decision bias to report target presence cannot apply. Thus, the effect of alpha amplitude lateralization just before stimulus onset, and thus before deliberation of the response, on comparative contrast judgments argues in favor of a bias at the subjective perceptual level, whereby elevated firing rates within the more excitable hemisphere are interpreted as if the stimulus had a higher contrast. This interpretation is in line with the finding that pre-stimulus alpha amplitude affects the strength of stimulus-evoked neuronal responses at the level of primary visual cortex (Iemi et al., 2019). Our comparative contrast discrimination task was adapted from studies on the effect of spatial attention on contrast appearance that used brief, non-informative cues just before stimulus onset for capturing attention. Numerous studies have demonstrated that stimuli presented at the cued location are judged as more contrasted compared to non-cued locations (Carrasco et al., 2004; Carrasco and Barbot, 2019). Thus, the conditions compared in the original paradigm (stimuli ipsilateral or contralateral to cue) are distinguished by an external experimental manipulation that is salient to the participants. By contrast, we did not use spatial cues in the contrast discrimination task and conditions were distinguished only by a spontaneous, internal physiological process (stimuli ipsilateral or contralateral to the more excitable hemisphere). What gave rise to these spontaneous shifts of excitability in the first place? Voluntary spatial cues induce an alpha amplitude decrement in the contralateral relative to the ipsilateral hemisphere, indicating greater excitability in the task-relevant hemisphere and greater inhibition in the task-irrelevant hemisphere (Sauseng et al., 2005; Thut et al., 2006; Foxe and Snyder, 2011) and such lateralization can also occur spontaneously in the absence of external cues, driven only by internal fluctuations in the deployment of spatial attention (Bengson et al., 2014; Boncompte et al., 2016). Moreover, we compared the spontaneous pre-stimulus lateralization patterns that predicted participants’ contrast judgments to lateralization patterns induced with conventional spatial cueing and found that these patterns were highly similar (Figure 2 and 3). Thus, a plausible speculation is that the spontaneous patterns of alpha-band lateralization observed in our contrast discrimination task reflects spontaneous shifts of spatial attention.

The absence of any external cues in our contrast discrimination task is also relevant vis-a-vis a criticism that has been leveled against studies on attention-induced effects on contrast appearance. Specifically, several studies have found that the effect of exogenous cues on comparative contrast judgments is strongest when the two stimuli are of similar contrast (Schneider and Komlos, 2008; Itthipuripat et al., 2019), raising the possibility that when participants are required to choose between two stimuli that look indistinguishable, cued stimuli might appear more salient or compelling to report (Schneider and Malik, 2021). This would constitute a bias at the decision level, not at the perceptual level. In fact, we also found that the effect of pre-stimulus alpha-band lateralization of contrast judgments was strongest when the objective contrast difference was minimal (Figure 4B). However, while participants are certainly aware of external cues, making them a potential confound in the decision process, there is no evidence that participants are similarly aware of internal, moment-to-moment hemispheric lateralization of excitability. Instead, it is more plausible that lateralization effects were strongest at minimal contrast differences because judgments are the least determined by the objective stimulus, such that subtle differences in excitability can tip the scales in favor of the stimulus represented in the more excitable hemisphere.

In addition to this within-subjects relationship between pre-stimulus alpha-band lateralization and contrast judgments, we found a strong relationship across subjects between individuals’ overall lateralization pattern and their general tendencies to judge one hemifield has having higher contrast (Figure 5B). Specifically, participants with a general idiosyncratic bias to judge, say, the left stimulus as having higher contrast showed stronger left-ward lateralization on trials when the left stimulus was judged as more contrasted, compared to less biased participants. This result suggests that the large inter-individual variability in subjective contrast perception is partially rooted in the same patterns of cortical pre-stimulus excitation and inhibition that also bias trial-by-trial perceptual experience. This finding nicely extends a set of previous research connecting individual biases with pre-stimulus, spontaneous alpha rhythm (Grabot and Kayser, 2020), by showing that local patterns of alpha increase and decrease not only influence the decision of participants at the single trial level, but that participants with different bias direction and strength show differential pre-stimulus patterns congruent with their own biases. This result is of outstanding interest, because it suggests one possible mechanism at the basis of the great intersubject variability in perception.

Our regression analysis showed, in addition to the effect of lateralization on contrast appearance, and the effect of bilateral pre-stimulus alpha amplitude on objective accuracy in the contrast judgment task: stronger amplitude was related to worse performance. This result was unexpected because it conflicts with recent studies showing that pre-stimulus alpha amplitude affects criterion, not accuracy in visual detection tasks (Limbach and Corballis, 2016; Iemi et al., 2017; Iemi and Busch, 2018), and does not affect performance in discrimination tasks (Samaha et al., 2017; Benwell et al., 2017; Samaha et al., 2020b; Benwell et al., 2021). Given that states of high alpha amplitude indicate low arousal (Johnston et al., 2020), diminishing the quality of stimulus representations (Zhou et al., 2021), one possibility is that the effect we observed was merely a by-product of fatigue, which in turn caused the effect on accuracy. However, note that the spatially specific effect of lateralized alpha amplitude on contrast perception in the contralateral hemifield cannot be explained by unspecific effects of fatigue.

## Conclusion

We demonstrate that the amplitude of pre-stimulus alpha oscillations alters subjective contrast appearance, such that perceived contrast is amplified for stimuli represented in the cortical hemisphere with weaker amplitude, and thus stronger neuronal excitability. These findings show that spontaneous oscillations affect not only decision making processes, but can alter our phenomenological perception of the visual world.

## Methods

### Participants

Forty-six participants took part in the experiment. Two of them were excluded because of performance below 2 SD from the population mean in the contrast discrimination task and 4 were excluded for noisy EEG signal. This left us with 40 participants (F=26, mean age=23 years, age range = 18-34 years, 34 right handed), the datasample size that had been pre-registered. All participants were compensated for participation with course credit or money (8 EUR/h). The study was approved by the ethics committee of the faculty of Psychology and Sports Science, University of Münster. All participants gave their written consent to participate.

### Apparatus

Recordings took place in a dimly-lit, sound-proof cabin. Participants placed their heads on a chin-rest and could adjust the height of the table to be seated comfortably. Stimuli were generated using Matlab 2019a (www.mathworks.com) and the Psychophysics Toolbox 3 (Brainard, 1997; Kleiner et al., 2007). The experiment was controlled via a computer running Xubuntu 16.04, equipped with an Intel Core i5–3330 CPU, a 2 GB Nvidia GeForce GTX 760 GPU, and 8 GB RAM. The experiment was displayed on a 24” Viewpixx/EEG LCD Monitor with 120 Hz refresh rate, 1 ms pixel response time, 95% luminance uniformity, and 1920*1080 pixels resolution (www.vpixx.com). Distance between participant eyes and the monitor was approximately 86 cm.

### Stimuli and experimental procedure

All stimuli were presented on a gray background (52.24 cd/*m*^2^). Each trial started with a fixation point, a small circle (diameter .1 dva) surrounded by 4 diagonal sectors of a bigger circle (diameter .6 dva) of a dark grey (6.5 cd/*m*^2^). Simultaneously with the fixation point, two placeholders of the same color were displayed, with an edge of 6.5 dva indicating the area of subsequent appearance of the gabors, and were centered at +-8.88 dva on the horizontal and at −4.6 dva on the vertical meridian, where all coordinates are referred to the center of the screen. Hence the placeholders and the stimuli appeared at 10 dva of eccentricity, in the lower visual field (VF). The fixation and the placeholders remained on the screen for the whole duration of the trial. After a variable SOA (uniformly sampled between 2100-3300 ms), two gabor patches (diameter = 6 dva; spatial frequency = 2 cpd) appeared for 41.67 ms (5 frames) in the positions delimited by the placeholders.

In the contrast discrimination task the gabor patches both had vertical orientations.. One stimulus, (standard) had a Michelson contrast of .2, whereas the other (test) could be in the range [−.45, .45] log10 units around the standard stimulus contrast. The contrast of the test stimulus was determined on a trial by trial basis by a Bayesian Adaptive procedure implemented ad hoc for the present experiment (see next paragraph). After the stimuli disappearance the participants had to report in which hemifield (left or right) the stimulus with the highest contrast had appeared.

In the orientation discrimination task, 1200 ms before stimuli appearance one of the lower sections (left or right) of the fixation point was increased in luminance (78.39 cd/*m*^2^) for 200 ms, indicating the position to be attended for reporting the orientation of the stimulus target. The stimuli consisted of 2 gabor patches which had the same Michelson contrast (.2) of the standard stimulus as defined before, but an orientation of ±45° with respect to the vertical axis. After the stimuli disappearance the participants had to report the orientation (left, counterclockwise; right, clockwise) of the cued stimulus. For both tasks participants were invited to provide their responses with their dominant hand by pressing left or right arrow keystroke on a standard German layout keyboard.

Throughout the whole experiment, each participant completed 768 trials, consisting of 192 trials for orientation discrimination and and 576 trials for contrast discrimination task. Of the latter, half were showing the test stimulus on the left hemifield and the other half on the right hemifield. The proportion of trials for each task (and for each hemifield probed) was maintained constant in the 8 blocks of 96 trials each. The order of the trial sequence was randomized. Every 24 trials, a summary feedback on performance was provided to the participant: such feedback was given collapsing together the orientation and contrast discrimination task, but was not taking into account those trials in which the choice was arbitrary (i.e. the standard and the test stimulus had the same contrast), since no definition of “correct response” could apply.

### Bayesian Adaptive procedure

In order to adapt the contrast of the stimuli on-line during the experiment, we implemented a custom Bayesian adaptive procedure in MATLAB following the guidelines detailed by Baek and coworkers (Baek et al., 2016) and by Watson (Watson, 2017). First we defined a model that could proficiently describe the probability of response toward the test stimulus. We individuated this model in a modified version of the logistic PMF as follows:

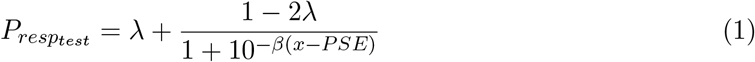

Where *x* is the contrast of the test stimulus compared to the standard, *λ* is the asymptotic value, *β* describes he slope of the PMF and *PSE* is the Point of Subjective Equality. It is possible to notice that in the current formulation *λ* describes both the low and the upper asymptotic value: this choice was made to stabilize the contrast probing on both the upper and the lower sides of the sigmoid in equal measure. Furthermore, during the online estimation, the *λ* value was set constant, in order to limit the amount of trials used to probe the extrema of the PMF.

In order to implement the adaptive procedure we defined a two-dimensional *θ* = (*PSE, β*) parameter space that represents all potential observable PMFs for each combination of parameters and a one dimensional stimulus search space over a range of possible contrasts for the test stimulus. Regarding the parameter space, values for *PSE* were selected in a range between [-.3313, .3313], leading to 41 equally spaced values on a logarithmical scale. 51 equally spaced values for *β* were selected in a range between [1.22, 10.88], whereas *λ*, as anticipated, was set constant at a value of .02. The ranges for the *PSE* and *β* parameters were chosen in order to encompass .99 confidence intervals of the parameter distributions obtained by fitting the model on pilot data. The contrast values were defined as 31 equally spaced log units in a range between [-.45, .45] around the standard contrast (log_10_(.2)). The procedure was implemented separately for the two visual fields.

### Online fixation control

Eye-movements were monitored using a desktop-mounted Eyelink 1000+ infrared based eye-tracking system (www.sr-research.com) set to 500 Hz sampling rate (binocular). The eye-tracker was (re-)calibrated using a nine-point calibration grid at default locations. Participants were required to keep their gaze on the fixation symbol throughout the trial, until the end of stimulus presentation. To ensure steady fixation and to avoid ocular or preferential encoding of one of the targets, a trial was aborted and repeated at the end of the respective block whenever participants blinked or gaze was outside of a 2.5° radius around the fixation symbol. The eye tracker was recalibrated at the start of each block and whenever participants lost fixation for more of three consecutive times, causing the restart of the trial.

### EEG recording and preprocessing

EEG was recorded with a Biosemi Active Two EEG system with 65 Ag/AgCl electrodes (www.biosemi.nl), set to 1024 Hz sampling rate. Sixty-four electrodes were arranged in a custom made montage with equidistant placement (“Easycap M34”; www.easycap.de), which extended to more inferior areas over the occipital lobe than the conventional 10–20 system (Oostenveld and Praamstra, 2001). An additional external electrode was placed below the left eye. EEG data were preprocessed using Matlab R2018a (www.mathworks.com) and the EEGlab toolbox (Delorme and Makeig, 2004), with the Cleanline (Mullen, 2012) extension. Data were resampled to 512 Hz, high-pass filtered at 1 Hz, low-pass filtered at 70 Hz, and notch-filtered between 48-52 Hz. Subsequently, data were rereferenced to a robust reference (Bigdely-Shamlo et al., 2015). Vertical and horizontal electrooculograms were derived from two electrodes above and below the left eye, and two electrodes at the lateral canthi of both eyes, respectively. Continuous data was segmented into epochs from 2000 ms before target onset to 500 ms after stimulus onset. To automatically clean the EEG signal from noisy segments, we applied a two step procedure. As a first step, in order to select artifacts coming from single electrodes in single trials we z-scored the EEG data for each trial along the channel dimension, and interpolated single segments exceeding an absolute z-value of 6. When a channel was interpolated for more than the 15% of the trials, then it was interpolated for all the trials. As a second step, epochs with irregular artifacts were automatically detected and rejected using a combination of threshold criterion, joint probability, or high power in higher frequency bands (40-70 Hz) indicating muscular noise. When the amount of data either interpolated or rejected exceeded the threshold of 15%, the data from a subject was discarded, as we preregistered. On average, the 7.454% ± 2.409 (mean±SD) of the data was either interpolated or discarded. Remaining artifacts were corrected using Independent Component Analysis (ICA). After ICA all electrodes were rereferenced to Common Average.

### EEG Data Analysis: pre-stimulus difference in timefrequency analysis

To explore the prestimulus timecourse and evaluate the range of frequencies involved in cue-induced attentional allocation and in the choice of a participant in the contrast discrimination task we performed a wavelet transform in fieldtrip (Oostenveld et al., 2011). The analysis was performed for frequencies ranging from 3 to 40 Hz, each wavelet of 3 cycles long, and sampling every 1 Hz and every 15.6 ms.

### EEG Data Analysis: pre-stimulus difference in instantaneous amplitude

We performed a pre-stimulus spectral analysis to elucidate the mechanisms underlying the decoding accuracy in both experimental conditions. In order to exclude any potential leak of poststimulus activity in the pre-stimulus window, and to limit edge artifacts due to filtering and to the Hilbert transform, for each trial we applied the following steps:

1. Selection of time window of interest [-500; 0] ms from stimulus onset.
2. Expansion of the signal by mirroring both tails for the first/last 195 ms.
3. Application of bandpass filter in the frequency range [8-13] Hz.
4. Application of Hilbert transform, and extraction of instantaneous amplitude by taking the absolute value of the complex signal.
5. Selection of the original portion of the signal.

We baselined the signal by subtracting, for each timepoint and each trial, the average amplitude of all channels. Next, for each experimental condition (orientation or contrast discrimination), we averaged together the trials belonging to “left” or “right” categories (cue direction or response, respectively for the task), and computed the difference for all channels. In order to compute lateralization, we selected a subset of 5 occipital channels for each side (left vs right, see Figure **??**), and we computed the difference between the averages of left and right channels. We were in this way able to obtain, for each single participant and each experimental condition the timecourse of alpha lateralization, between comparisons of interest. Most importantly, positive sign of this timecourse indicated increased in alpha contralateral to participants’ response, or cued location. Statistical significance was evaluated with one-sample t-test against 0. To correct for multiple comparisons, we performed a cluster permutation test, where clusters were defined by contiguity in the time domain: for 10000 repetitions, we multiplied a random participants subset by −1, we computed the cluster statistic by selecting all the contiguous points with *p* < .05, creating distribution of randomly generated cluster statistics (Samaha et al., 2017). Then we computed the empirical cumulative distribution function for the random distribution and we individuated the probability value for the cluster statistic obtained with the non-shuffled data. We rejected *H*_0_ when the cluster statistic was exceeding the 95° percentile of permutations.

### EEG Data Analysis: Similarity between cue-induced and spontaneous lateralization

Topographical similarity between cue-induced and spontaneous lateralization was computed via matrix multiplication. So, for each participant, let *W* be the *N_channels_* × *M_timepoints_* produced by the difference between left and right cue, and *X* the matrix (of same size), produced by the difference in the response, similarity was computed as:

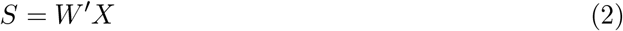

Where ′ indicates matrix transpose. This leads to a *M_timepoints_* × *M_timepoints_* matrix which shows how similar are the two patterns across the whole time window.

Notably we evaluated the difference in similarity between trials were the perceptual uncertainty was higher vs those trials in which the perceptual uncertainty was lower. We defined perceptual uncertainty based on the inverse of the PMF in Eq. 1 as follows:

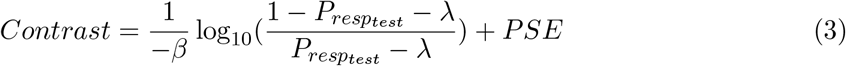

In this way, we were able to select the trials whose contrast difference between test and standard stimulus was belonging to the range of contrasts around the threshold (*P* = .5 ± .25, “high uncertainty”) and the ones outside this range, defined as “low uncertainty”. For each of the two groups of trials, within each participant, we computed the difference between left and right cue, and the difference in response as described before. Based on Eq. 2, we evaluated whether the “high uncertainty” trials, *X_high_* were showing a stronger similarity to the cue induced lateralization than the “low uncertainty” *X_low_*:

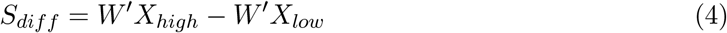

At the group level, we tested whether this difference was different from 0 with one sampled ttests on the whole similarity matrix. We corrected for multiple comparisons via a cluster permutation approach analogous to the one defined before, but applied this time on the similarity matrix instead of a time series.

### EEG Data Analysis: Logistic regression analysis

Preprocessing steps premiliminary to the logistic regression analysis include 3 additional steps, so we expand here the procedure already descripted above:

1. Selection of time window of interest [-500; 0] ms from stimulus onset.
2. Expansion of the signal by mirroring both tails for the first/last 195 ms.
3. Application of bandpass filter in the frequency range [8-13] Hz.
4. Downsampling to 128 Hz, to decrease computation time.
5. Application of Hilbert transform, and extraction of instantaneous amplitude by taking the absolute value of the complex signal.
6. Selection of the original portion of the signal.
7. Rank scoring across trials to mitigate the effect of outliers.
8. Z scoring across trials in order to obtain normalized *β* scores from the model.

Importantly, the last step (Z scoring) was applied to all model’s regressors. The model was defined as a typical logistic regression model in the form:

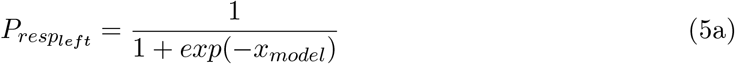

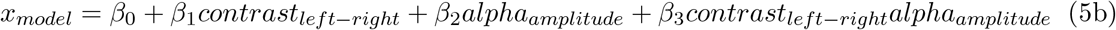

Our model consisted of 4 different regressors as described in Eq. 5b: *β*_0_, indicating the overall tendency to respond toward one or the other hemifield independently from the actual contrast of the stimuli and underlying brain activity, *β*_1_, the weight that the physical contrast difference between stimuli had in determining the outcome of the decision process, *β*_2_ the weight of prestimulus alpha amplitude in *β*_3_, as the weight for the interaction between physical contrast difference and pre-stimulus alpha amplitude. Logistic regression coefficients were computed via iterative reweighted least square (Bishop, 2006), for each combination of electrode and timepoint for the whole pre-stimulus window. The application of the model yielded hence for each participant a *N_timepoints_* × *M_timepoints_* × 4 regressors for each subject.

At the group level, we tested whether the weights for the pre-stimulus alpha amplitude and the interaction between alpha amplitude and physical contrast difference were significantly different from 0. For this purpose, we first ran a one-sample t-test against 0, for each channel, timepoint and regressor. We defined clusters by selecting those t-values exceeding the critical threshold (*α* = .05) on both tails. From each cluster we obtained a cluster statististic by summing together all the t-values belonging to the cluster, i.e. characterized by contiguity in space (electrodes) and time. To correct for multiple comparisons, for 1000 repetitions, we multiplied a random participants subset by −1, and repeated the abovementioned steps in order to obtain a distribution of cluster statistics under the null hypothesis. We rejected the null hypothesis if a cluster statistic was below the 2.5° or above the 97.5° percentiles of the permutations.

### EEG Data Analysis: Correlation between spontaneous lateralization and individual Points of Subjective Equality (PSE)

PSEs were computed according to Eq. 3, by defining *P* = .5, for each participant and visual field. We aggregated the PSEs between visual fields as follows:

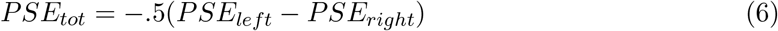

where *PSE_left_* and *PSE_right_* are the perceptual thresholds estimated for the left and right VF, and *PSE_tot_* the aggregated one. The sign correction was applied in order to maintain consistency across the results, so that negative values indicate bias toward the right hemifield, and viceversa. As a second step, we followed the same procedure used to evaluate the spontaneous patterns of lateralization already described before (see *pre-stimulus difference in instantaneous amplitude*), consisting of the difference between left and right occipital channels in the contrast between left and right response. This yielded a timecourse of strength and directionality of alpha lateralization, that we correlated with the PSE of the participants, for each timepoint in the pre-stimulus window. This yielded a timeseries of correlations spanning the 500 ms before stimulus onset. We tested for multiple comparison by first identifying the cluster showing contiguous timepoints with *p* < .05 (two-tailed), and summing up the *r* values for all the point in the cluster to obtain a relative cluster statistic. Then, for 1000 times, we shuffled the vector of PSEs and repeated the steps before, in order to to obtain a distribution of cluster statistics under the null hypothesis. We rejected the null hypothesis if a cluster statistic was below the 2.5° or above the 97.5° percentiles of the permutations.

## Data Accessibility

The data and the code used to produce the figures and compute the statistics will be made public upon acceptance of the current manuscript.

## Acknowledgments

This work was supported by a grant from the German Research Foundation (DFG; BU2400/9-1). We thank Teresa Berther, Luca Michels and Louisa Henkels for help with data acquisition and Jason Samaha for helpful comments at an early stage of data analysis.

## Notes

### Competing Interest Statement

The authors have declared no competing interest.

